# Tbet expression by Tregs is needed to protect against Th1-mediated immunopathology during *Toxoplasma* infection in mice

**DOI:** 10.1101/2021.09.03.458860

**Authors:** Jordan Warunek, Richard M. Jin, Sarah J. Blair, Matthew Garis, Brandon Marzullo, Elizabeth A. Wohlfert

**Author notes:** Corresponding Author: Elizabeth A. Wohlfert.

## Abstract

*T. gondii* infection has proven to be an ideal model to understand the delicate balance between protective immunity and immune-mediated pathology during infection. Lethal infection causes a collapse of Tregs mediated by loss of IL-2, and conversion of Tregs to IFNγ producing cells. Importantly, these Tregs highly express the Th1 transcription factor Tbet. To determine the role of Tbet in Tregs, we infected *Tbx21*^*f/f*^-Foxp3^YFPCre^ and control Foxp3^YFPCre^ mice with the type II strain of *T. gondii*, ME49. The majority of *Tbx21*^*f/f*^-Foxp3^YFPCre^ mice succumb to a non-lethal acute infection. Notably, parasite burden is comparable between *Tbx21*^*f/f*^-Foxp3^YFPCre^ and Foxp3^YFPCre^ control mice. We found that *Tbx21*^*f/f*^-Foxp3^YFPCre^ mice have significantly higher serum levels of proinflammatory cytokines IFNγ and TNFα, suggestive of a heightened immune response. To test if CD4^+^ T cells were driving immunopathology, we treated *Tbx21*^*f/f*^-Foxp3^YFPCre^ mice with anti-CD4 depleting antibody and partially rescued these mice. Broad spectrum antibiotic treatment also improved survival, demonstrating a role for commensal flora in immunopathology in *Tbx21*^*f/f*^-Foxp3^YFPCre^ mice. RNA-seq analysis reinforced that Tbet regulates several key cellular pathways, including chromosome segregation, cytokine receptor activity and cell cycle progression, that help to maintain fitness in Tregs during Th1 responses. Taken together, our data shows an important role for Tbet in Tregs in preventing lethal immunopathology during *Toxoplasma gondii* infection, further highlighting the protective role of Treg plasticity to self and microbiota.

## Introduction

The sustained presence of regulatory T cells (Tregs) is vital to maintaining immunologic homeostasis and required for host survival as evidenced by the emergence of autoimmune diseases when Treg suppressive function is disrupted (1, 2). Tregs not only regulate autoimmune responses but they also modulate immune responses to invading pathogens and defective Treg functionality has been implicated in a spectrum of pathologies (1, 3). As the immunological landscape rapidly changes during an infection, Tregs must adapt to the inflammatory environment. This adaptation, termed Treg plasticity, endows them with certain features of T helper cells that enhance survival and functionality of Tregs in diverse inflammatory conditions (4–11). For example, the acquisition of certain Th1 effector properties aids in enhancing suppression during Th1 inflammation and the upregulation of CXCR3 confers precise homing and accumulation at these sites, however not all of the acquired phenotypes are fully understood (12–15). Tregs in T helper 1 cells (Th1) polarized inflammatory environments will also upregulate Tbet, the lineage-specifying transcription factor for Th1 cells (4–11). During both acute and chronic *T. gondii* infection in mice, Tbet-expressing Tregs can be found throughout the host (12, 16–20), including the small intestinal lamina propria, brain, muscle, spleen and lymph nodes (11, 12, 16–20). Notably, Tregs in skeletal muscle express equivalent levels of Tbet compared to CD4^+^ effector T cells (16). Yet, its functional role in Tregs during *T. gondii* infections had not been explored.

Th1-like Tregs (Tbet^+^ Tregs) are a highly stable population of Tregs, suggesting their persistence is advantageous to the host if they possess a suppressive function and their specificity is towards self-antigen (14). Our lab has previously shown that, during chronic *T. gondii* infections, Th1-like Tregs are perpetrators of muscle damage by promoting macrophages to maintain proinflammatory M1 properties instead of tissue regenerative M2 phenotypes (16). These disparate findings in immunoregulatory processes led us to question whether Tbet expression in Tregs was essential for host response and control of *T. gondii*. This is important because there is also debate on the role of Tbet in Tregs within the inflamed gastrointestinal tract and if Tbet is needed for Treg function (21, 13, 22). Taken together, these ambiguities led us to question whether Tbet expression in Tregs is required for host response and control of *T. gondii* infection.

To address this question, we infected Treg-specific Tbet deficient conditional mice, *Tbx21*^*f/f*^-Foxp3^YFPCre^, and control Foxp3^YFPCre^ mice with the type II strain ME49. In our attempt to understand if Tbet was driving the pathology in chronically infected muscle, we discovered that Tbet is required for survival of the host during acute infection. Interestingly, while Tbet expression by Tregs was required for survival of acutely infected mice, it did not influence host control of the parasite. We found that both CD4^+^ T cells and the commensal flora partially drove the immunopathology observed in the *Tbx21*^*f/f*^-Foxp3^YFPCre^ mice during acute infection with *T. gondii*. RNA sequencing on sorted splenic Tregs from day 10 infected *Tbx21*^*f/f*^-Foxp3^YFPCre^ and Foxp3^YFPCre^ mice highlighted several cellular pathways regulated by *Tbx21* in Th1-Tregs. Among these include pathways related to chromosome segregation, cytokine receptor activity and cell cycle progression. Taken together this reveals a critical role for Tbet in mediating the fitness of the Treg response to infection induced dysbiosis, and remains a vital determinant of host survival during oral *T. gondii* infection.

## Materials and Methods

### Mice

B6.129(Cg)-^*Foxp3tm4(YFP/icre)Ayr*^*/*J (Foxp3^YFPCre^) and B6.129-*Tbx21*^*tm2Srnr*^/J (*Tbx21*^*fl/fl*^) were purchased from Jackson Laboratories (Bar Harbor, ME) and crossed to generate *Tbx21*^*f/f*^-Foxp3^YFPCre^ mice. All experimental mice were raised in SPF conditions and were sex and age matched and used at 8 to 10 weeks old at the time of infection. Both males and females were used in experiments. All procedures involving mice were reviewed and approved by the Institutional Animal Care and Use Committee at the University at Buffalo.

### Isolation of tissue lymphocytes from organs

Spleen and mesenteric lymphocytes were harvested by passing through Falcon 70μm cell strainers to create single suspensions in 2% media (RPMI with 2% FBS, 25 mM HEPES, 0.1% β-mercaptoethanol, 1% penicillin-streptomycin, 1% L-Glutamine). Red blood cells were lysed as needed on ice for 3 minutes and resuspended in 10% media (RPMI with 10% FBS, 25 mM HEPES, 0.1% β-mercaptoethanol, 1% penicillin-streptomycin, 1% L-Glutamine).

Liver was collected and finely minced in digestion media (RPMI, 1% penicillin-streptomycin, 1mM sodium pyruvate, 0.1% β-mercaptoethanol, 25mM HEPES, 150μg/ml DNase I [Sigma-Aldrich], 0.5mg/ml Liberase [Roche] and then slow spun at 50xg for 5 mins (11). Small intestinal lamina propria (SILp) were isolated as previously described (11). SILp was harvested, cut into 0.5 cm pieces and pre-digested in 3% Media with 5mM EDTA and 0.145mg/mL DTT for 20 minutes at 37° C. The lamina propria was isolated from flow through epithelium by shaking in 2mM EDTA, 25mM Hepes, and 1% penicillin-streptomycin media. The tissue was then finely minced in digestion media containing Liberase and DNase I and incubated for 30 minutes at 37°C degrees (11). Digested tissue was passed through a 70μM and then 40μM strainer. SILP single cell suspensions were resuspended in 10% media (RPMI with 10% FBS, 1% penicillin-streptomycin, 1mM sodium pyruvate, 0.1% β-mercaptoethanol, 25mM HEPES).

### Extracellular and Intracellular flow cytometric analysis of tissue lymphocytes

Single-cell suspensions were stained with HBSS containing extracellular surface antibodies and Live/Dead Fixable Blue Dead Cell Stain (Life Technologies). After extracellular staining for 25 minutes on ice, cells were fixed and permeabilized overnight (Intracellular Fixation and Permeabilization Buffer Set, Thermo Fisher eBioscience). Cells were then washed and stained with the eBioscience Permeabilization Buffer containing intracellular antibodies for 45 minutes on ice. Samples were washed and resuspended in flow cytometry buffer/FACS Buffer (PBS, 1% bovine serum albumin [Sigma-Aldrich], 2mM EDTA [Life Technologies]) for acquisition on a BD Fortessa SORP. Absolute numbers were derived from cell counts using a hemocytometer (Life Technologies) or CountBright Absolute Counting Beads (Invitrogen).

### *T. gondii*–specific tetramer staining

Single-cell suspensions of tissue lymphocytes were stained with allophycocyanin-conjugated MHC class II tetramers bound to *T. gondii* ME49 hypothetical antigenic peptide I-A(b) (AVEIHRPVPGTAPPS) obtained through the National Institute of Health Tetramer Facility (Atlanta, GA). Cells were incubated with the *T. gondii–*specific tetramer or CLIP tetramer for 1 hour at room temperature. After *T. gondii*-specific staining, cells were stained with extracellular and intracellular antibodies as described previously.

### Antibodies

Antibodies used in flow cytometric analysis were: anti-TCRβ-APC-Cy7 (BD Pharmigen, clone H57-597), anti-CD4-PE-Cy7 (BD Pharmigen, clone RM4-5), anti-CD4-V450 (clone RM4-5, BD Pharmingen), anti-CD8b-Percp-Cy5.5 (clone H35-17.2, BD Pharmingen), anti-CD11a-BV650 (clone 2D7, BD Pharmingen), anti-CD49D-BV421 (clone R1-2, BD Pharmingen), anti-Foxp3-FITC (eBioscience, clone FJK-16s), anti-Tbet-PE (eBioscience, clone eBio4B10), anti-Ki67-AF700 (BD Pharmingen, clone B56). Flow cytometry data was acquired using a BD LSRFortessa Cell Analyzer and analyzed using FlowJo version 10.0.8 (Tree Star, Ashland, OR).

### Detection of serum cytokines

Sera was isolated from mice either by cardiac puncture or collected by submandibular vein puncture at various time points. Assessment of the sera cytokines was performed by ELISA to detect IFNγ and TNFα (eBioscience) at 10 to 12 days post-infection.

### *Toxoplasma* infection

Brains were isolated from chronic *T. gondii* infected mice (30 to 60 days post-infection). The brain was homogenized in phosphate-buffered saline pH 7.2 (PBS) for oral infection (12). RFP-expressing ME49 cysts from brain homogenates were enumerated using a fluorescence microscope to detect RFP and counted in triplicate. Mice were orally gavaged with five ME49^RFP^ cysts in 200ul PBS. Mouse weight, activity and posture were scored every other day for the first 8 days of infection, monitored daily during acute infection at day 8 through day 14 and then weekly thereafter.

### Antibody depletion of CD4^+^ T cells *in vivo*

Groups of *Tbx21*^*f/f*^-Foxp3^YFPCre^ and Foxp3^YFPCre^ mice were treated with either 0.5mg of anti-mouse CD4, (clone GK1.5), or rat IgG2b isotype control anti-keyhole limpet hemocyanin, (clone LTF-2) (Bio X Cell’s InVivoMab). Mice were intraperitoneally injected with a depleting antibody or isotype control on days −5, −1, 0 pre-infection and on days 5, 14, 19 post-infection.

### Antibiotic Treatment

To deplete gut microbiota, groups of Foxp3^YFPCre^ and *Tbx21*^*f/f*^-Foxp3^YFPCre^ mice were given broad-spectrum antibiotics in their drinking water (23). The following antibiotics were added to autoclaved tap water: ampicillin (1 g/L), vancomycin (500 mg/L), neomycin sulfate (1 g/L), and metronidazole (1 g/L) (Sigma-aldrich, St. Louis, MO) and administered *ad libitum* using drinking bottles. A set of mice from each group were designated as untreated controls and received water supplied from the animal facility. Antibiotic treatment started two weeks prior to infection with *T. gondii* and was administered until three weeks post-infection, a total of five weeks of treatment.

### Analysis of bacterial 16S genes

Feces were collected prior to antibiotic water administration and at various time points thereafter. Fecal pellets were collected directly into microcentrifuge tubes and immediately placed on dry ice until storage in a −80 freezer. Bacterial DNA was isolated using the QIAamp Fast DNA Stool Mini Kit (Qiagen). Quantification of DNA was done using a nanodrop. qPCR was performed using the iTaq Universal SYBR Green Mastermix (BioRad) on a StepOne™ Real-Time PCR System. Bacterial detection was analyzed using amplification of the 16s ribosomal subunit of specific populations. The selected bacteria to be analyzed by qPCR were Eubacteria, Enterobacteriaceae, *Escherichia coli*, Bacteroides, Eubacterium rectale/Clostridium coccoides group (EREC) (24). Primer sequences are included in Table 1.

### Quantification of parasite burden

Mouse organ tissues were harvested and preserved in RNAlater (Invitrogen) for DNA isolation using a DNeasy Blood & Tissue Kit (Qiagen). DNA was quantified using a Nanodrop Spectrophotometer for quality and concentration. An amount of 200ng of various tissue DNA was used in PCR amplification of the *T. gondii* specific repetitive gene B1 (forward: 5′-TCCCCTCTGCTGGCGAAAAGT-3′, reverse: 5′-AGCGTTCGTGGTCAACTATCGATTG-3′). Parasite burden was calculated using a comparison of a standard curve of known *T. gondii* genomic DNA amplified with *B1* specific primers.

### Blood chemistry assays and complete blood count

Blood was collected by cardiac puncture after euthanasia from *T. gondii* infected mice. For chemistry analysis, the blood was placed in tubes without anticoagulants and placed on ice until analysis. UB LAF veterinary staff analyzed blood chemistry using an Element DC Veterinary Chemistry Analyzer (Heska). For complete blood counts, blood was placed on ice in tubes coated with EDTA and analyzed using an Element HT5 Veterinary Hematology Analyzer (Heska).

### RNA-Seq

Tregs were sort purified from *Tbx21*^*f/f*^-Foxp3^YFPCre^ and Foxp3^YFPCre^ mice 9 days after being infected with five ME49 cysts. Total RNA to be sequenced was extracted from 1×10^6^ sort purified Tregs using TRIzol (Thermo Fisher). RNA sequencing was performed and analyzed at the UB Genomics and Bioinformatics Core Facility using the R package DESeq2 to determine differentially expressed genes between Tregs from infected *Tbx21*^*f/f*^-Foxp3^YFPCre^ and Foxp3^YFPCre^. Gene ontology analysis was performed using the GOseq Bioconductor package version 1.42.0. GOseq performs Gene Ontology analysis, while addressing biases present in RNA-seq data that is not found using other techniques, namely that expected reads counts for a transcript is based on both the gene’s level of expression, and the length of the transcript. The Wallenius approximation is used by GOSeq to approximate GO category enrichment and calculate p-values for each GO category being over represented among genes that were differentially expressed (DESeq2 padj <= 0.05). These enrichment p-values were corrected for multiple testing using the Benjamini-Hochberg Procedure.

### Statistics

All statistics were generated using Graphpad Prism v6.0h.

## Results

### Tbet expression in Tregs is required for survival of acute T. gondii infection

We investigated the role of Tbet in Tregs during *Toxoplasma* by administering 5 ME49 *T. gondii* cysts via oral gavage to both Foxp3^YFPCre^ and *Tbx21*^*f/f*^-Foxp3^YFPCre^ mice. The acute phase of infection begins in the intestinal tract, and is marked by weight loss. We found a similar weight loss between Foxp3^YFPCre^ and *Tbx21*^*f/f*^-Foxp3^YFPCre^ infected mice (Figure 1A). However, *Tbx21*^*f/f*^-Foxp3^YFPCre^ infected mice succumbed to infection at a significantly higher rate (87%) than infected Foxp3^YFPCre^ mice, where the majority survived through acute infection (Figure 1B). We first asked if *Tbx21*^*f/f*^-Foxp3^YFPCre^ mice were succumbing to parasitemia because of an inability to control the infection. To this end, we harvested tissue from day 12 post-infected mice and used qPCR to quantify *T. gondii* DNA using the tandem repeat gene *B1*. No differences were observed in parasite burden between mice in the thymus, lung, heart and ileum. There was significantly less parasite in the brains of *Tbx21*^*f/f*^-Foxp3^YFPCre^ mice versus Foxp3^YFPCre^ controls (Figure 1C). These findings show that Tbet expression in Tregs is necessary for survival of acute *T. gondii* infection, and that it does not negatively impact immune control of *T. gondii* infection.

**Figure 1.**
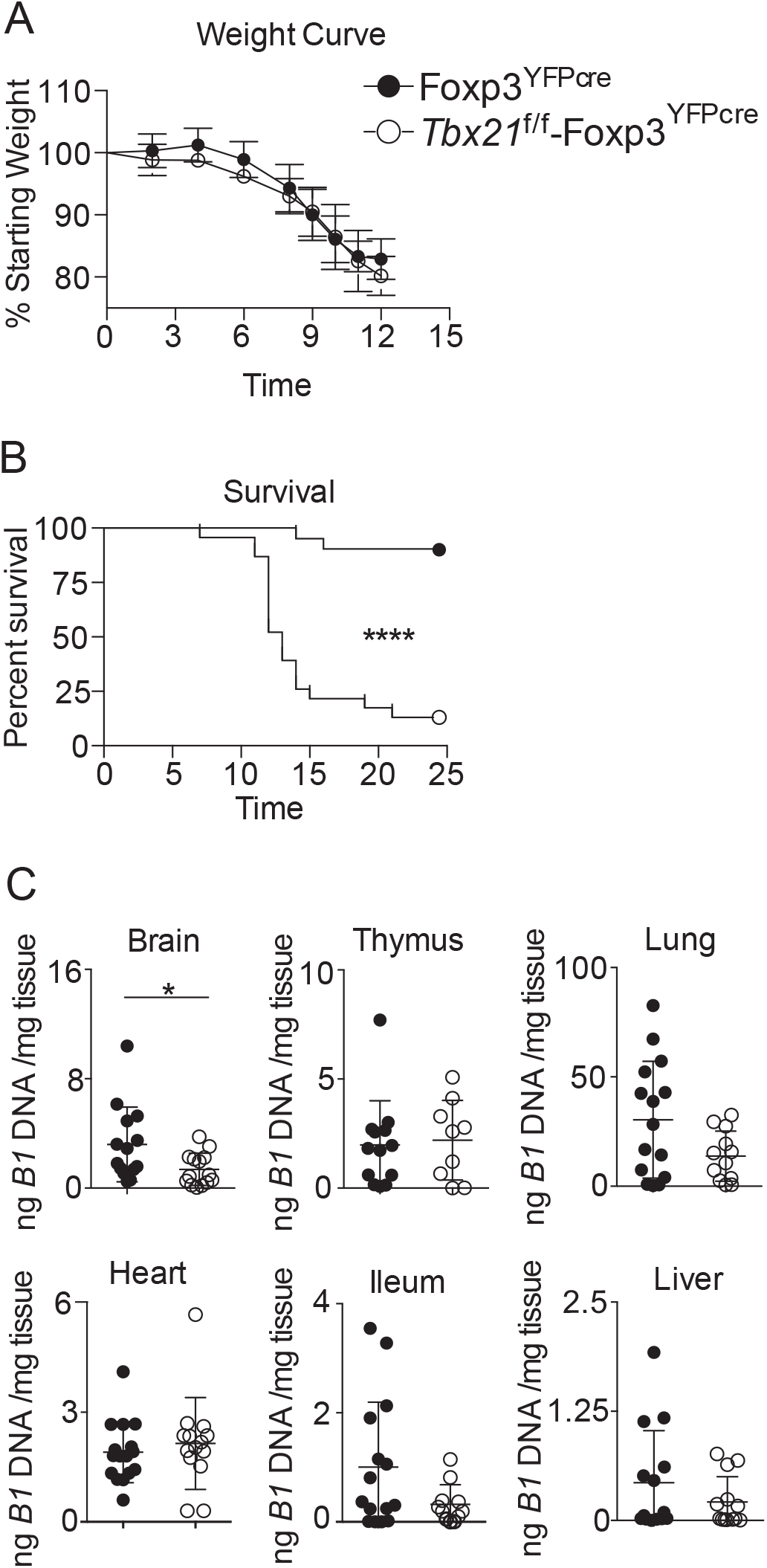
Tbet expression in Tregs is required for survival to *T. gondii* infection. Groups of Foxp3^YFPCre^ and *Tbx21*^f/f^-Foxp3^YFPcre^ mice were orally infected with five ME49^RFP^ *T. gondii* cysts. Tissues were isolated at 12 d postinfection and parasite burden was quantified. (**A**) Percent weight loss and (**B**) survival curve summary from four experiments using male and female mice. (**C**) Tissues from day 12 infected Foxp3^YFPCre^ and *Tbx21*^f/f^-Foxp3^YFPcre^ mice were harvested to extract gDNA and analyzed for parasite burden using *T. gondii-*specific gene, *B1*. Results are cumulative of four experiments with n ≥ 3 mice per group per experiment. A cumulative total of n ≥ 16 per group (A,B) and n ≥ 9 (C); error bars = SD. ^*^p<0.05, ^****^p<0.0001, Log-rank (Mantel-Cox) Test (B), Student *t* test (C).

### Tbx21^f/f^-Foxp3^YFPCre^ mice exhibit severe pathology during acute T. gondii infection

Since uncontrolled parasite growth was not observed in *Tbx21*^*f/f*^-Foxp3^YFPCre^mice, we turned to the host response to understand how Tbet expression in Tregs could promote host survival during acute infection. We analyzed biomarkers in serum associated with excessive inflammation to ascertain if acute organ failure was occurring in the absence of Treg-specific Tbet expression. A comprehensive metabolic panel was performed on blood freshly isolated from *Tbx21*^*f/f*^-Foxp3^YFPCre^ mice and Foxp3^YFPCre^ control mice day 10 post-infection. Significantly decreased levels of glucose and albumin were observed in the *Tbx21*^*f/f*^-Foxp3^YFPCre^ mice. In addition, significantly higher levels of inorganic phosphate, cholesterol and blood urea nitrogen were measured in the *Tbx21*^*f/f*^-Foxp3^YFPCre^ mice (Figure 2A). These differences in the serum may be tied to liver and kidney damage, however when measuring ALT/AST levels we found no significant differences between the two groups of mice (data not shown). In addition to blood metabolic panels, a complete blood count was performed to determine if hematopoiesis was differentially altered in *Tbx21*^*f/f*^-Foxp3^YFPCre^ mice compared to Foxp3^YFPCre^ mice during acute infection. Total leukocytes, lymphocytes, eosinophils, and neutrophils were not significantly different. However, a significantly lower frequency and number of monocytes were observed within the blood of *Tbx21*^*f/f*^-Foxp3^YFPCre^ (Figure 2B). This data suggests that in spite of the immune response controlling the infection, there was potentially systemic lethal immunopathology in *Tbx21*^*f/f*^-Foxp3^YFPCre^ mice.

**Figure 2.**
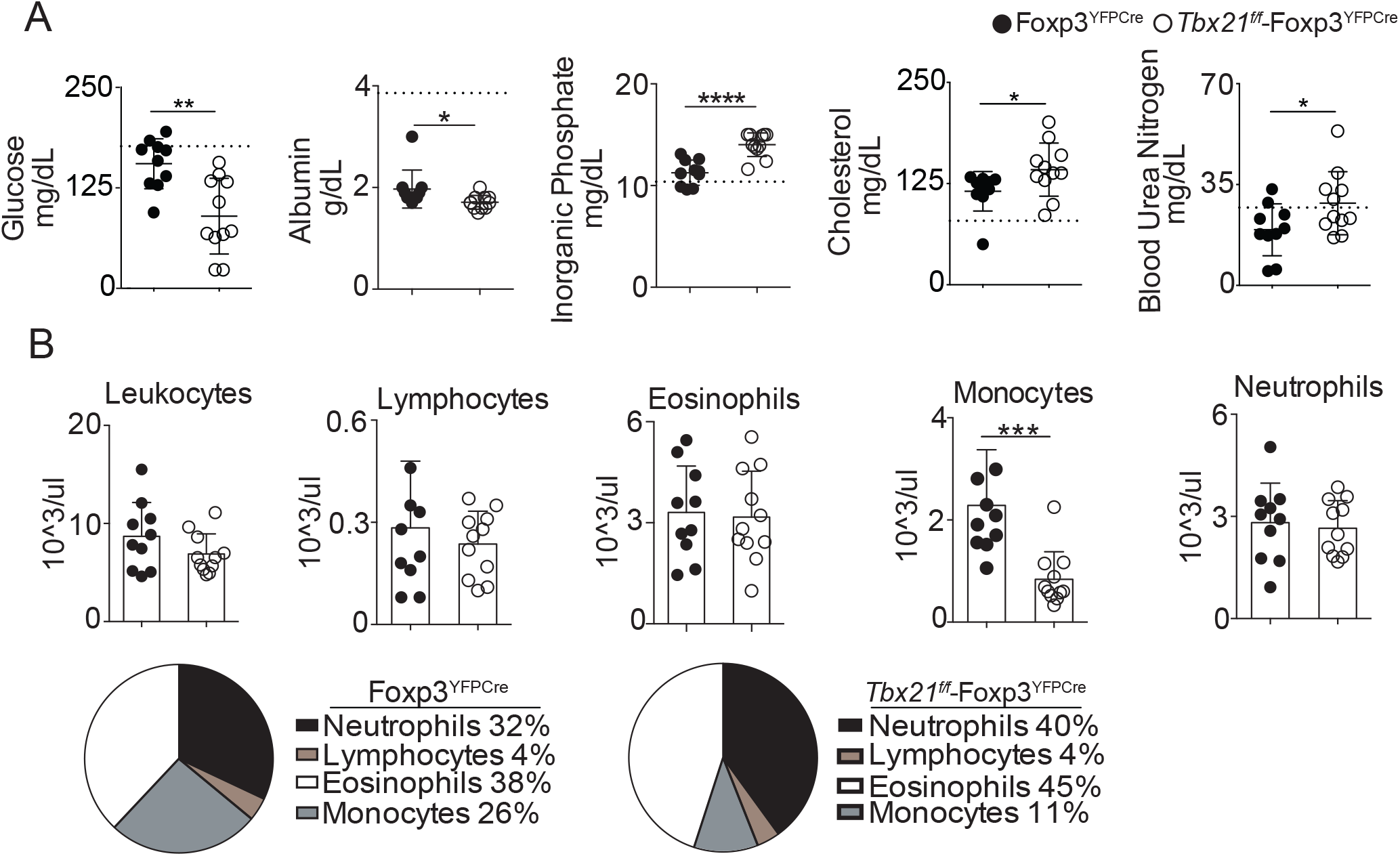
Acute *T. gondii* infection causes immunopathology and metabolic shift indicative of systemic damage in *Tbx21*^f/f^-Foxp3^YFPcre^ mice. Groups of Foxp3^YFPCre^ and *Tbx21*^f/f^-Foxp3^YFPcre^ mice were orally infected with five ME49^RFP^ *T. gondii* cysts. Mice were euthanized 10 d postinfection using CO_2_ and fresh blood was isolated by cardiac puncture to perform a comprehensive metabolic panel and complete blood count. (**A**) Blood chemistry assay was performed 10 d postinfection looking at liver and kidney function. Dashed lines are reference values for uninfected mice (**B**) Complete blood counts 10 d postinfection quantifying absolute numbers of leukocytes and (**C**) pie graphs of percent leukocytes in the blood. Results are cumulative of 2 experiments with n ≥ 5 per group per experiment. A cumulative total of n ≥ 10 per group (A-C). Error bars = SD. ^*^p<0.05,^**^p<.01, ^***^p<0.001,^****^p<0.0001, Student *t* test.

### *Treg-specific Tbet deficiency elevates systemic IFNγ and TNFα, but not T helper 1 populations during acute* T. gondii *infection*

We next examined the CD4^+^ T cell immune response in *T. gondii* infected *Tbx21*^*f/f*^-Foxp3^YFPCre^ mice compared with control Foxp3^YFPCre^ mice. Analysis of serum levels of IFNγ and TNFα, critical Th1 inflammatory cytokines that are important in host survival to *T. gondii* (25–29), were significantly increased in *Tbx21*^*f/f*^-Foxp3^YFPCre^ mice compared to Foxp3^YFPCre^ mice (Figure 3A). Next, CD4^+^ T cells were examined by FACS analysis for phenotype and absolute numbers in the spleen, mesenteric lymph nodes (mln), and small intestinal lamina propria (SILp). There was no difference in the absolute number of CD4^+^TCRβ^+^Foxp3^-^ T cells (Figure 3B), or their expression of Ki67^+^(Figure 3C). The frequency of Tbet in CD4^+^TCRβ^+^Foxp3^-^ T cells was similar in the mln and SILp but was reduced in the spleen in *Tbx21*^*f/f*^-Foxp3^YFPCre^ mice compared to Foxp3^YFPCre^ mice (Figure 3D). We assessed the binding of the AS15 tetramer, which identifies *T. gondii* antigen specific CD4^+^ T cells. We found no difference in the frequency or absolute number of AS15 tetramer positive CD4^+^ T cells in the mln and SILp but both were reduced in the spleen in *Tbx21*^*f/f*^-Foxp3^YFPCre^ mice compared to Foxp3^YFPCre^ mice (Figure 3E). These results suggest that while expression of Tbet by Tregs is important for the restraint of wide-spread, fatal inflammation, it is largely dispensable for limiting the size and activity of the Th1 cell pool and cell parasite-specific CD4+ T cell responses in the gastrointestinal tract during acute infection.

**Figure 3.**
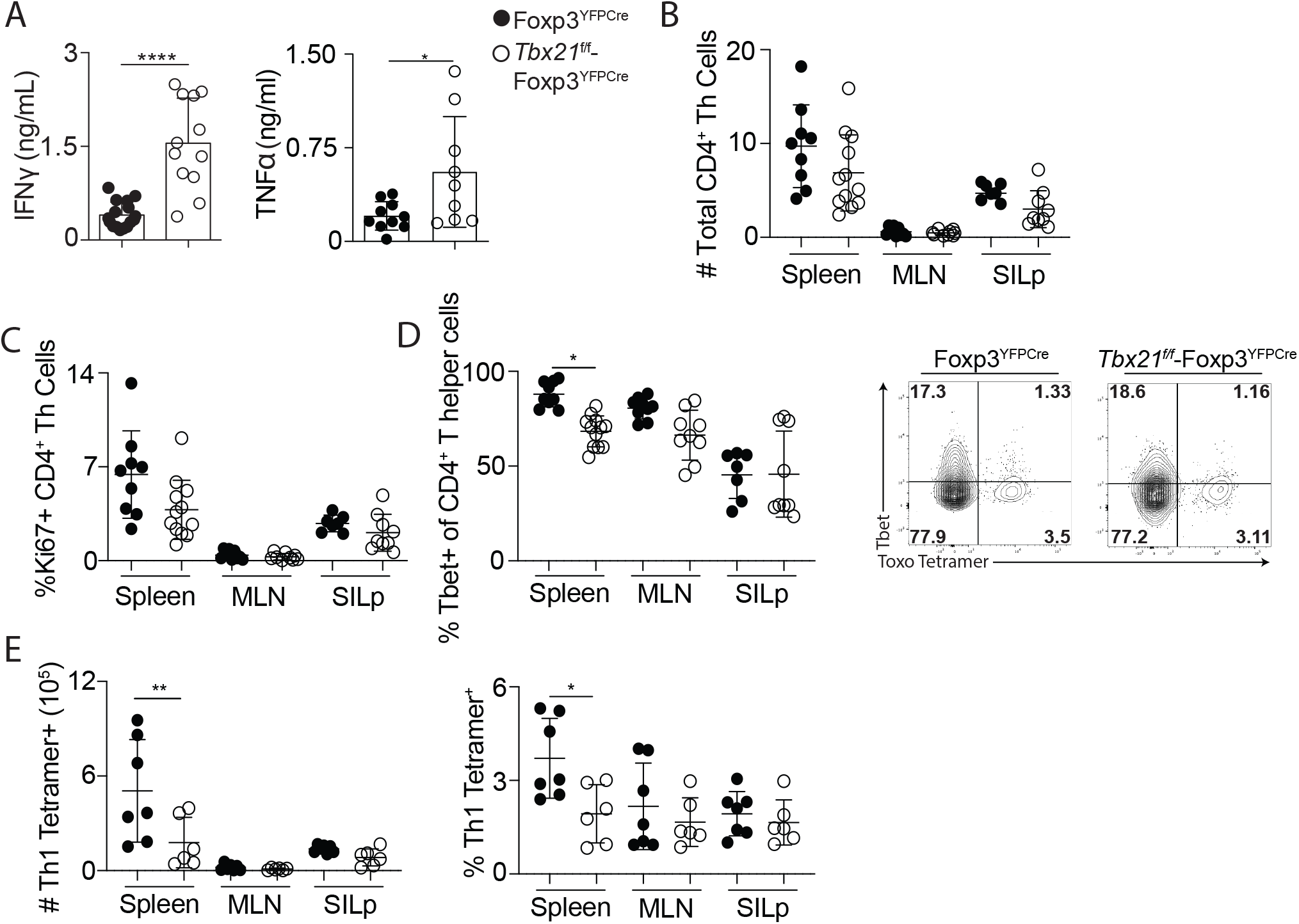
T helper cells recognize *T. gondii* in similar fashion during acute infection. Groups of Foxp3^YFPCre^ and *Tbx21*^f/f^-Foxp3^YFPcre^ mice were orally infected with five ME49^RFP^ *T. gondii* cysts. Lymphocytes from the spleen, mesenteric lymph nodes (mln), and small intestinal lamina propria (SILp) were isolated on 12-14 d postinfection. CD4+ T helper cells were stained and analyzed using flow cytometry. (**A**) ELISA was performed to quantify IFNγ (left) and TNFα (right) in sera 10-12 d postinfection. (**B**) Quantification of absolute total number of CD4^+^ T helper cells, gated on live TCRβ^+^CD4^+^Foxp3^-^ cells. (**C**) Percentages of Ki67^+^ or Tbet^+^ T helper cells in spleen, mln, and SILp. (**D**) Representative FACs plots of *T. gondii* tetramer staining, gated on live TCRβ^+^CD4^+^Foxp3^-^ looking at T helper coexpression of Tbet and tetramer in SILp. (**E**) Absolute (left) and frequency (right) quantification of Th1 *T. gondii* tetramer staining, gated on live TCRβ^+^CD4+Foxp3^-^Tbet^+^ Tetramer^+^. n ≥ 3 per group, cumulative of 3 experiments. A cumulative total of n ≥ 6 per group. Error bars = SD. ^**^p < 0.01, ^*^p < 0.05, ANOVA with Tukey multiple comparison test.

### *Tbet is required for the proliferation and accumulation of Tregs in the SILp during acute* T. gondii *infection*

We next examined CD4^+^Foxp3^+^ Tregs in the small intestinal lamina propria, mesenteric lymph nodes, and spleens of infected mice by FACS analysis. We found there was a reduced frequency of Tregs in *Tbx21*^*f/f*^-Foxp3^YFPCre^ mice compared to Foxp3^YFPCre^ mice in the mesenteric lymph node, however no difference in the total number of Tregs was observed (Figure 4A). Tregs in the SILp were reduced in absolute numbers (Figure 4A left panel). Tregs from the SILp had reduced frequency of the proliferation marker Ki67 (Figure 4B left panel) and reduced absolute number of Ki67-expressing Tregs (Figure 4B right panel) in *Tbx21*^*f/f*^-Foxp3^YFPCre^ mice compared to Foxp3^YFPCre^ mice. As expected, Tregs from *Tbx21*^*f/f*^-Foxp3^YFPCre^ mice compared to Foxp3^YFPCre^ mice expressed significantly less Tbet (Figure 4C).

**Figure 4.**
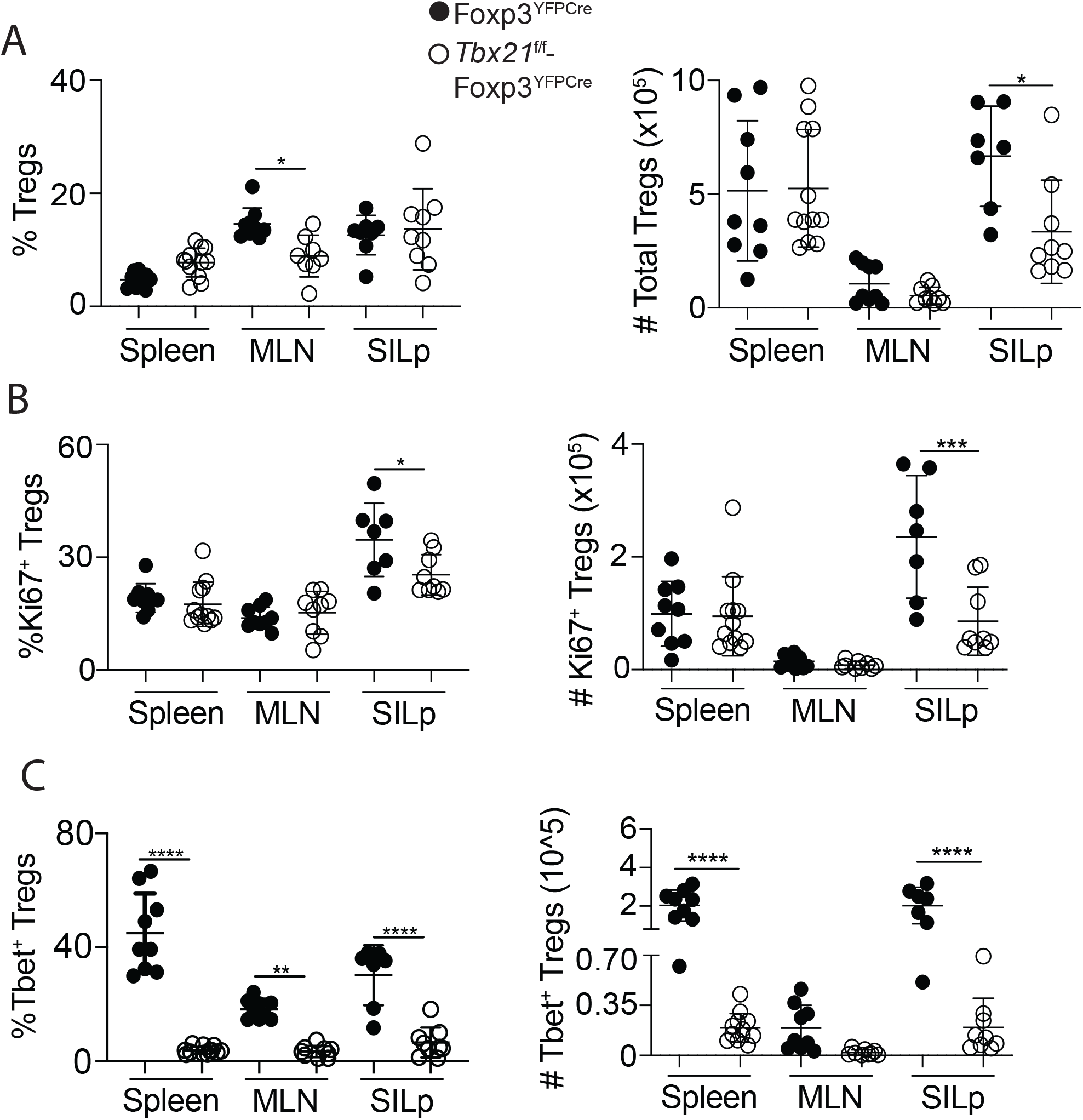
Comparison between Tregs during acute infection. (**A**) Quantification of Treg frequency (left) and absolute (right), gated on live TCRβ^+^CD4^+^Foxp3^+^ in the spleen, mesenteric lymph node (mln) and small intestinal lamina propria (SILp) on day 12-14 dpi. (**B**) Frequency (left) and absolute (right) of Ki67+ Tregs. (**C**) Frequency (left) and absolute (right) of Tbet^+^ Tregs. Results are cumulative of at least 3 experiments with n ≥ 3 per group. A cumulative total of n ≥ 7 per group. Error bars = SD. ^*^p < 0.05, ^**^p < 0.01, ^****^p<0.0001, ANOVA with Tukey multiple comparison test (C-D).

Since the increased mortality and immunopathology in the *Tbx21*^*f/f*^-Foxp3^YFPCre^ mice were potentially indicative of inflammation associated with multi-organ failure, we also examined the T cell response in the liver. We found a small but significant decrease in the frequency of CD4^+^TCRβ^+^Foxp3^-^ T helper cells, but no difference in the total numbers. We find no difference in the number of antigen experienced CD4^+^ Th1 cells (CD11a^+^CD49d^+^), their expression of Tbet or the proliferative marker, Ki67 between *Tbx21*^*f/f*^-Foxp3^YFPCre^ mice and Foxp3^YFPCre^ mice (Figure 5A). Analysis of CD8^+^TCRβ^+^ in the liver showed no differences between the two groups of mice in frequency, numbers, activation status and proliferative capacity (Figure 5B). We examined the same parameters on Tregs in the liver of infected mice. Notably, we found that while small intestinal Tregs from *Tbx21*^*f/f*^-Foxp3^YFPCre^ mice were reduced in both numbers and proliferation, there was a significant increase of both frequency and number of Tregs in the liver (Figure 5C). This data suggests that Tbet is needed for Treg accumulation at primary sites of acute infection, but secondary sites may not rely on Tbet for Treg accumulation.

**Figure 5.**
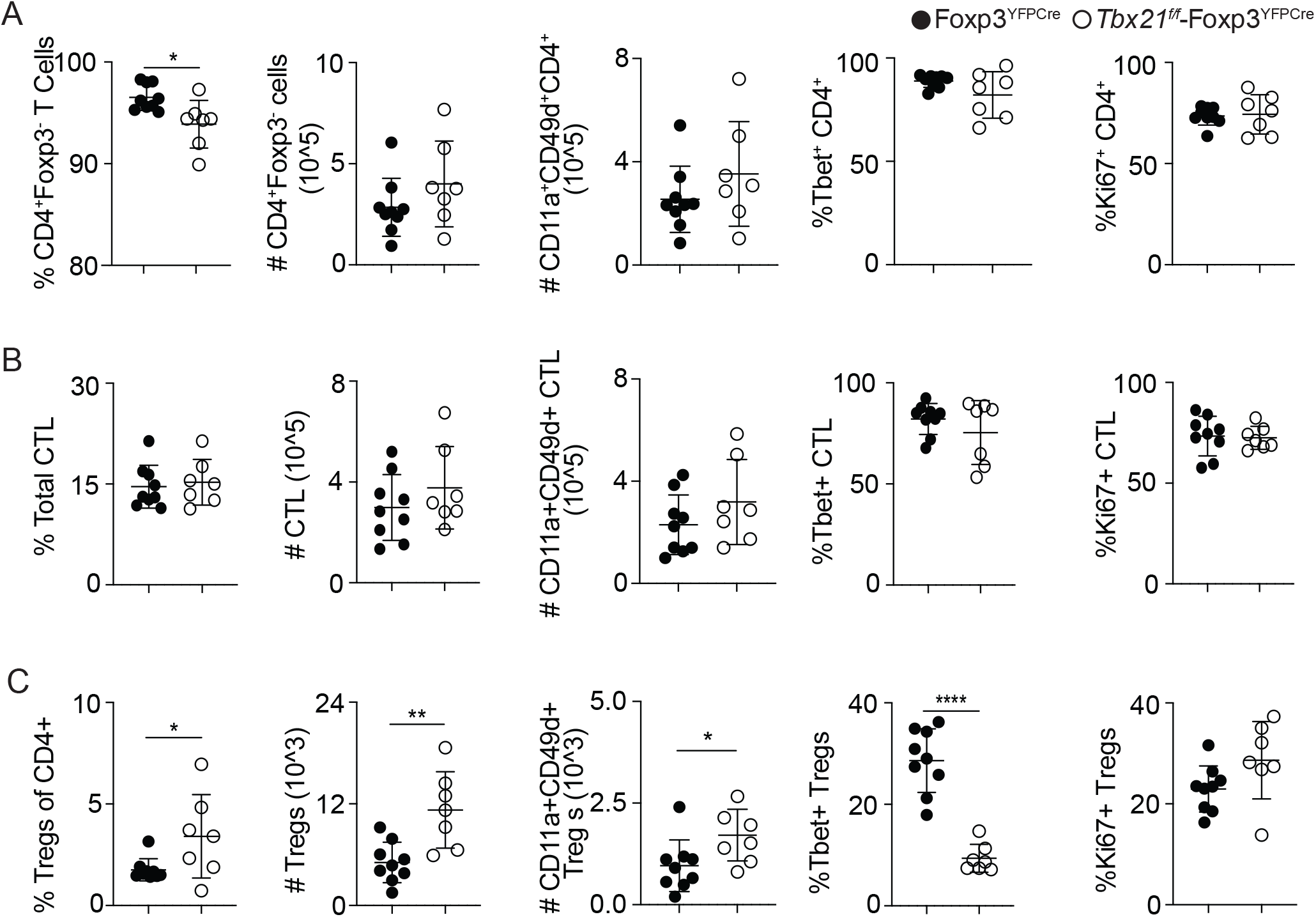
Differences in Tregs but not CD4 or CD8 T cells in the liver during acute infection. (A) Quantification of T helper cells gated on live CD4^+^TCRb^+^Foxp3^-^ in the liver on day 12 dpi. (B) or CD8^+^TCRb^+^ or (C) Tregs, CD4^+^TCRb^+^Foxp3^+^. Results are cumulative of 2 experiments with n ≥ 3 in each group. Error bars = SD. ^*^p < 0.05,^**^p < 0.01,^****^p < 0.0001, Student’s t-Test.

### *CD4*^*+*^ *T cells participate in the enhanced immunopathology from acute* T. gondii *infection in Tbx21*^*f/f*^*-Foxp3*^*YFPCre*^ *mice*

We next asked if the morbidity and mortality observed in the *Tbx21*^*f/f*^-Foxp3^YFPCre^ mice was T cell mediated, which is observed in other strains of mice where immune regulation is altered (30, 31). The increased levels of serum IFNγ and TNFα in (Figure 3A) prompted us to deplete CD4 T cells using the anti-CD4 monoclonal antibody, GK1.5. Mice were I.P. injected with 0.5mg/mL anti-CD4 depleting or isotype antibody (clone LTF-2) before and during infection (30). Infected Foxp3^YFPCre^ mice that received anti-CD4 antibody had similar survival compared with isotype control treated infected Foxp3^YFPCre^ mice. However, *Tbx21*^*f/f*^-Foxp3^YFPCre^ mice that received anti-CD4 depleting antibodies were partially rescued from *T. gondii* infection compared with isotype control treated *Tbx21*^*f/f*^-Foxp3^YFPCre^ mice (Figure 6A). We found no difference between Foxp3^YFPCre^ or *Tbx21*^*f/f*^-Foxp3^YFPCre^ mice treated with either isotype control or anti-CD4 antibodies and parasite burden (Figure 6B). We next determined if there was a reduction in the serum levels of IFNγ or TNFα in Foxp3^YFPCre^ mice compared with *Tbx21*^*f/f*^-Foxp3^YFPCre^ mice treated with isotype control or anti-CD4 antibodies. As observed previously, *Tbx21*^*f/f*^-Foxp3YFPCre mice had significantly higher serum levels of IFNγ as well as heightened TNFα levels compared to Foxp3YFPCre mice. However, no marked differences in the levels of these cytokines were seen upon CD4 depletion (Figure 6C). The increased levels of IFNγ and TNFα in *Tbx21*^*f/f*^-Foxp3^YFPCre^ mice, despite anti-CD4 treatment, highlights the importance of fully functional Tregs to not only control T helper cells, but also other arms of immunity. The partial rescue of CD4-depleted *Tbx21*^*f/f*^-Foxp3^YFPCre^ mice implicates a role for CD4^+^ T cells in driving lethal immunopathology when Tregs cannot express Tbet during *T. gondii* infection.

**Figure 6.**
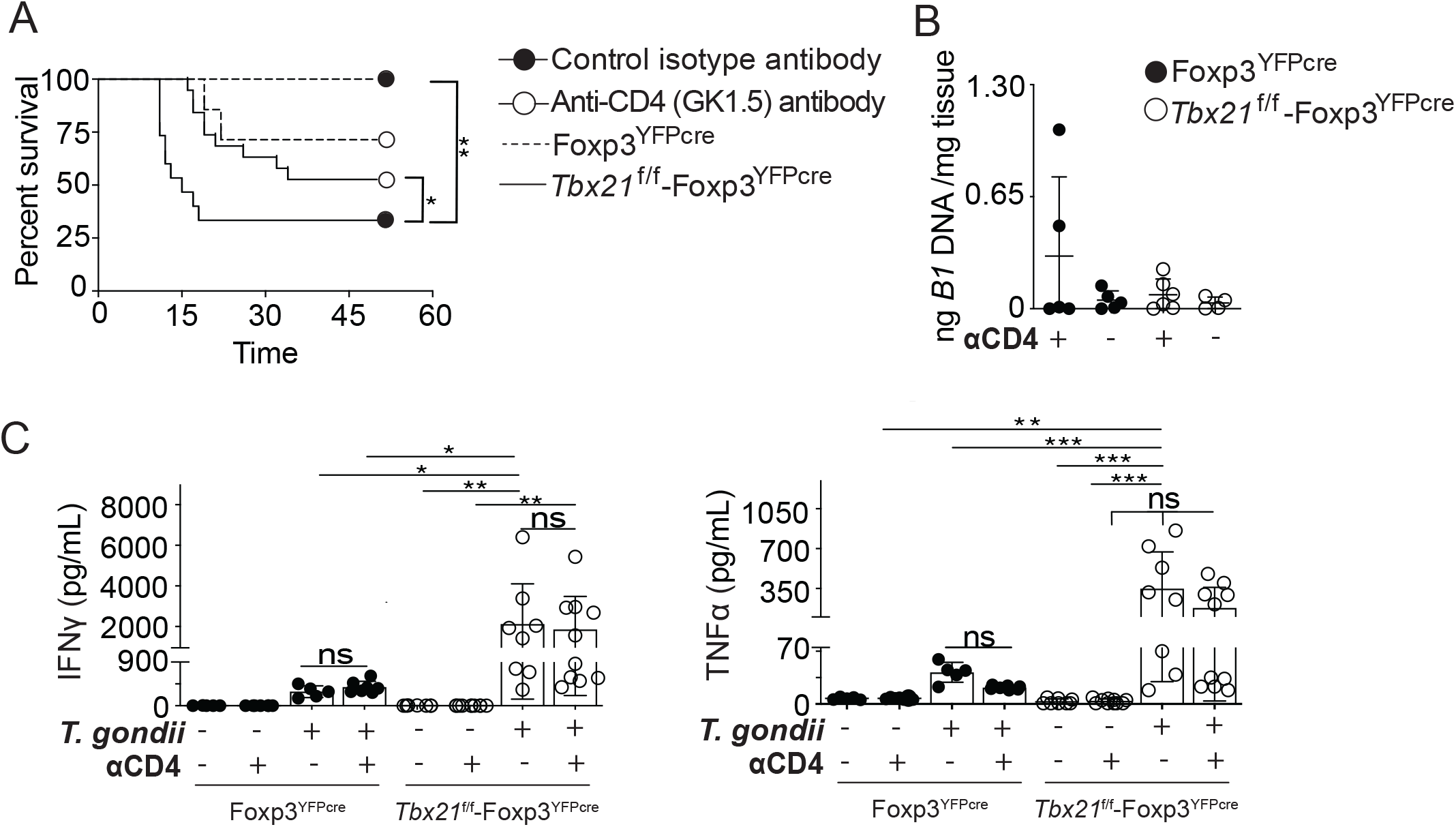
CD4 depletion partially rescues *Tbx21*^f/f^-Foxp3^YFPcre^ mice from succumbing to *T. gondii* infection. Groups of Foxp3^YFPCre^ and *Tbx21*^f/f^-Foxp3^YFPcre^ mice were I.P. injected with anti-CD4 depleting antibody or isotype control before and after infection on days −5, −1, 0, +5, +14, +19. All mice were orally infected with five ME49^RFP^ *T. gondii* cysts. (**A**) Survival curve from mice orally infected with five ME49^RFP^ cysts and I.P. injected with 0.5 mg/mL anti-CD4 depleting antibody or isotype control. (**B**) Parasite burden in the brain at day 42 postinfection, quantified by DNA extraction and using qPCR to amplify *T. gondii B1* gene. (**C**) Cytometric Bead Array to quantify IFNγ (left) and TNFα (right) serum cytokine levels on 9-10 d postinfection. Survival curve is cumulative of four experiments with n ≥ 4 per group per experiment. CBA is cumulative of two experiments with n ≥ 3 per group per experiment. A cumulative total of n ≥ 5 Foxp3^YFPCre^ and n ≥ 15 *Tbx21*^f/f^-Foxp3^YFPcre^ mice were used in the survival curve (A). A cumulative total of n ≥ 4 mice per group (B-C). Error bars = SD. ^*^p < 0.05,^**^p < 0.01 ^***^p<0.001, ^****^p<0.0001, Log-rank (Mantel-Cox) Test (A) ANOVA with Tukey multiple-comparison test (B,C)

### *Broad spectrum antibiotics ameliorates immunopathology from acute* T. gondii *infection in* Tbx21^f/f^-*Foxp3*^*YFPCre*^ *mice*

Mice infected perorally with a high dose *T. gondii* succumbed to severe ileitis and small intestine immunopathology through Th1 associated proinflammatory cytokines (32–34). This acute Th1 immune response causes a shift in the gut commensals communities and can contribute to the morbundity and mortality of mice during acute *T. gondii* in genetically susceptible hosts (32, 35, 36). Dysbiosis is attributed to specifically accumulated populations of the Gram-negative bacteria *E. Coli*, Bacteroides spp and other proinflammatory associated bacteria (37, 38). The use of broad spectrum antibiotics to deplete commensal gut flora rescues mice from systemic immunopathology during the acute stages of *T. gondii* infection (37, 38). To test if the lethal immunopathology in *Tbx21*^*f/f*^-Foxp3^YFPCre^ mice was mediated by an over exuberant immune response to commensal flora, we administered water supplemented with broad spectrum antibiotics to both strains of mice 14 days prior to infection with *T. gondii*. Mice received water without antibiotics as controls. Mice were orally gavaged with 5 ME49 cysts and received continued antibiotic treatment until 21 days post-infection. Foxp3^YFPCre^ mice showed no difference in survival between control water and antibiotic water (Figure 7A). The *Tbx21*^*f/f*^-Foxp3^YFPCre^ cohort that was administered antibiotic water survived significantly longer than the *Tbx21*^*f/f*^-Foxp3^YFPCre^ controls. Of note, 50% of the mice receiving antibiotics in the *Tbx21*^*f/f*^-Foxp3^YFPCre^ group survived until day 65 post-infection (Figure 7A). We found no difference between Foxp3^YFPCre^ mice mice that received control water or antibiotic water. *Tbx21*^*f/f*^-Foxp3^YFPCre^ mice that received antibiotic water had reduced cyst burden compared to Foxp3^YFPCre^ mice treated with antibiotic water (Figure 7B).

**Figure 7.**
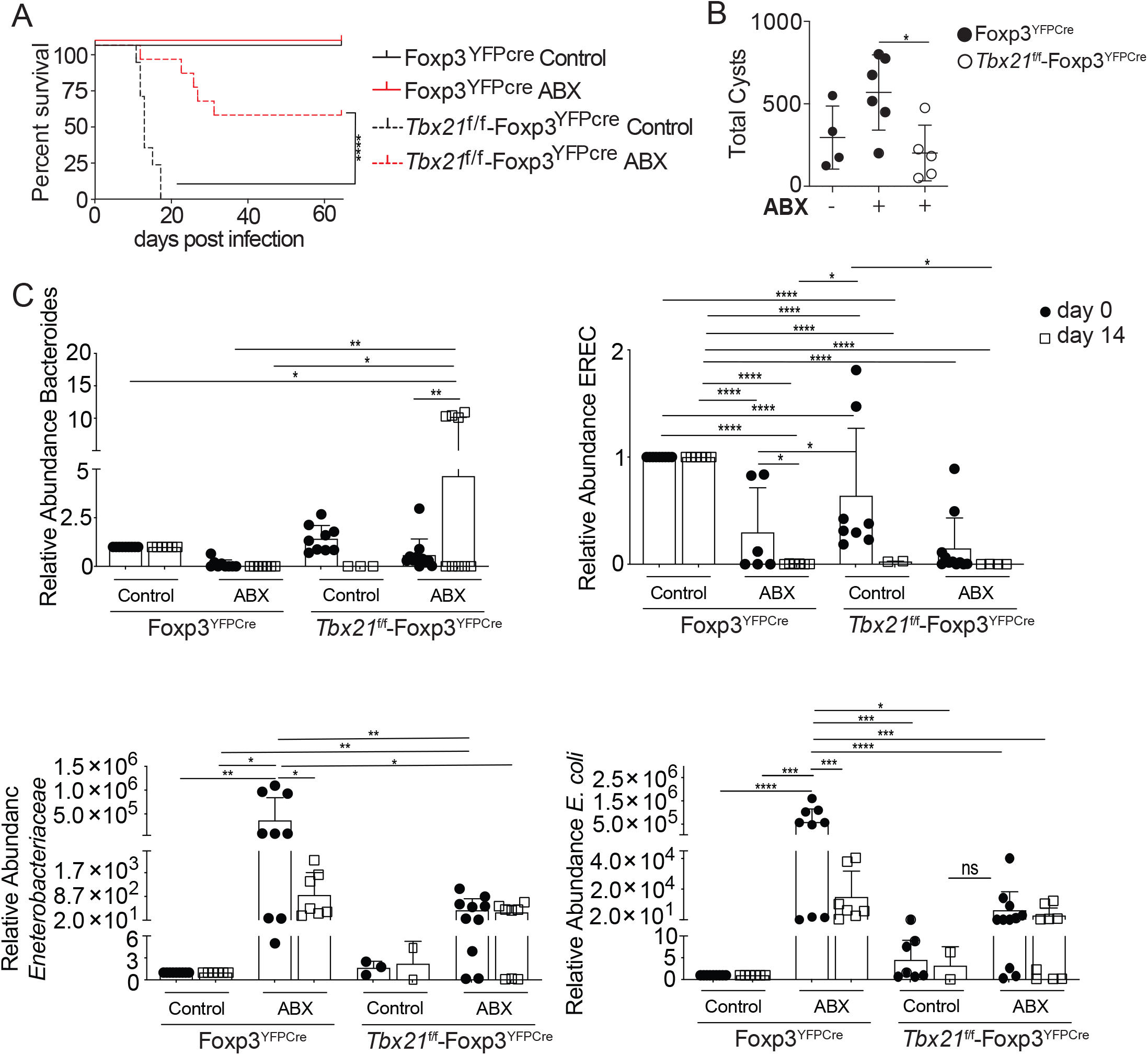
Commensals contribute to the mortality of *Tbx21*^f/f^-Foxp3^YFPcre^ during infection. Foxp3^YFPCre^ and *Tbx21*^f/f^-Foxp3^YFPcre^ were treated with broad spectrum antibiotic water containing ampicillin 1g/L, neomycin 1g/L, metronidazole 1g/L, vancomycin 500 mg/L and Splenda 3g/L. antibiotic water was administered in drinking bottles *ad libitum* for 2 weeks prior and 3 weeks after infection with *T. gondii*. Fecal samples were collected weekly to analyze bacterial populations. (**A**) Survival curve of antibiotic treated and water control mice after being infected with five ME49^RFP^ cysts with a 65 day postinfection endpoint. (**B**) ME49^RFP^ cyst counts from the brain homogenate on day 65 postinfection. (**C**) qPCR on DNA to semiquantitatively detect the 16s gene in fecal pellets collected for each of the following bacterial populations: Bacteroides, Eubacterium rectale/Clostridium coccoides (EREC), Enterobacteriaceae, and *Escherichia coli* (*E*.*coli*). Foxp3^YFPcre^ untreated control mice were used as reference. Results are cumulative of three experiments with n ≥ 3 per group per experiment. A cumulative total of n ≥ 9 per group (A), n ≥ 4 (B) and n ≥ 6 except for Tbet water controls (C) Error bars = SD. ^****^p < 0.0001, ^*^p < 0.05, Mantel-cox log-rank test (A), ANOVA with Tukey multiple-comparison test (C).

Since antibiotic treatment partially rescued the *Tbx21*^*f/f*^-Foxp3^YFPCre^ mice, we wanted to investigate the microbiota landscape and how subsequent dysbiosis caused by infection was altered between antibiotic treated and non-treated groups. To verify if the overall bacterial load was depleted, we compared 16s levels from before antibiotic treatment to the levels on day 7 post infection (Supplemental Figure 1). After 3 weeks of antibiotic administration in both Foxp3^YFPCre^ and *Tbx21*^*f/f*^-Foxp3^YFPCre^ groups, 16s was significantly lower on the day 7 post infection timepoint. This result confirms that bacterial load in the antibiotic treated groups was decreased as expected. To parse out the specific changes in bacterial populations, qPCR was performed using primers targeting unique 16s rRNA subunit gene in DNA to discriminate global bacterial changes of Enterobacteriacae family, *Bacteroides* family, Eubacterium rectale/Clostridium coccoides group (EREC), and *E. coli* species (24). Quantifying the changes of the microbiota was achieved by comparing pre-infection at day 0 and 14 days post-infection. Unfortunately, due to the nature of *Tbx21*^*f/f*^-Foxp3^YFPCre^ mice succumbing to infection, we lacked comparisons to the *Tbx21*^*f/f*^-Foxp3^YFPCre^ water control group at day 14. Therefore, *Tbx21*^*f/f*^-Foxp3^YFPCre^ antibiotic water treated mice were compared with the Foxp3^YFPCre^ water control group. We first asked if there were baseline differences between the four groups of mice prior to infection at day 0, after mice had been pre-treated for 2-weeks with antibiotic water. Our results show a significant outgrowth of Enterobacteriaceae and a species of that family, *E. coli* in both antibiotic and treated groups. This increase was present in both antibiotic groups, but the Foxp3^YFPCre^ antibiotic group had an even greater increase in Enterobacteriaceae and *E. coli* when compared to the *Tbx21*^*f/f*^-Foxp3^YFPCre^ antibiotic-treated samples. While Enterobacteriacae and *E. coli* were easily quantified in both antibiotic groups, we had difficulty detecting it in the *Tbx21*^*f/f*^-Foxp3^YFPCre^ control group. Another distinction between groups was that the EREC population was detected at day 0 in the *Tbx21*^*f/f*^-Foxp3^YFPCre^ controls but was depleted in their counterpart antibiotic groups.

Next, we examined the differences in bacterial populations at day 14 to assess changes within the infected mice. Our results showed a significant increase in the Bacteroides group that was specific to the *Tbx21*^*f/f*^-Foxp3^YFPCre^ antibiotics group on day 14 of infection. The Bacteroides population was also absent within the *Tbx21*^*f/f*^-Foxp3^YFPCre^ antibiotic group prior to infection. Finally, we examined the consequences of antibiotic treatment on dysbiosis and any changes between Foxp3^YFPCre^ control group at day 14 and antibiotic treated mice. There were significant decreases of EREC in both antibiotic groups at day 14 when compared to Foxp3^YFPCre^ controls. Also, the distinct Bacteroides group at day 14 in the antibiotic *Tbx21*^*f/f*^-Foxp3^YFPCre^ cohort was absent in the infected Foxp3^YFPCre^ controls. Lastly, Enterobacteriacae and *E. coli* increase during infection was significantly higher in the Foxp3^YFPCre^ antibiotic group compared to the controls. Although there was an increase of Enterobacteriaceae and *E. coli* at day 14 in the *Tbx21*^*f/f*^-Foxp3^YFPCre^ antibiotic group, this was not significantly different from day 14 Foxp3^YFPCre^ controls or antibiotics. The increase in Enterobacteriaceae and *E. coli* in the *Tbx21*^*f/f*^-Foxp3^YFPCre^ antibiotic cohort was significantly different at day 0, but not at day 14. The increase that was observed prior to infection could be due to antibiotics depleting the majority of bacteria and supports *E. coli* opportunistically blooming in comparison to other resident commensals (37, 38). Together, our data supports the idea that immunopathology induced by dysbiosis is exacerbated in *Tbx21*^*f/f*^-Foxp3^YFPCre^ mice infected with *T. gondii*.

### Tbet promotes Treg cell cycle progression and fitness

We sought to investigate global genetic changes in Tregs during infection and how Tbet influences Treg gene expression. We isolated total RNA from FACS sorted splenic Tregs (CD4^+^YFP^+^) 10 days post-infection from *Tbx21*^*f/f*^-Foxp3^YFPCre^ and Foxp3^YFPCre^ mice for RNA sequencing. There were 399 genes that were upregulated and 576 that were downregulated in Tregs from *Tbx21*^*f/f*^-Foxp3^YFPCre^ compared with Foxp3^YFPCre^ mice (Figure 8A). There are several differentially expressed genes of note that are downregulated between in infected *Tbx21*^*f/f*^-Foxp3^YFPCre^ compared to Foxp3^YFPCre^ sort purified Tregs, including *Tbx21, Miat, Nkg7, Gzmk, Ccl4, Ccl5*, and *Hip1*. Differentially expressed genes that are upregulated in Tbet deficient Tregs include *Hpgds, Il1rn, Il1r1 Areg* and *Penk* (Figure 8B). GOSeq analysis showed that several biological processes associated with cell cycle progression and mitotic processes are significantly enriched (Figure 8C), which is consistent with the reduction in proliferation and accumulation of Tbet-deficient Tregs in our previously reported data (Figure 3) and (5). Taken together, Tbet drives the expression of several aspects of fitness, in addition to homing molecules (5), that promote the ability of Tregs to prevent the immunopathology caused by dysbiosis during acute *T. gondii* infection.

**Figure 8.**
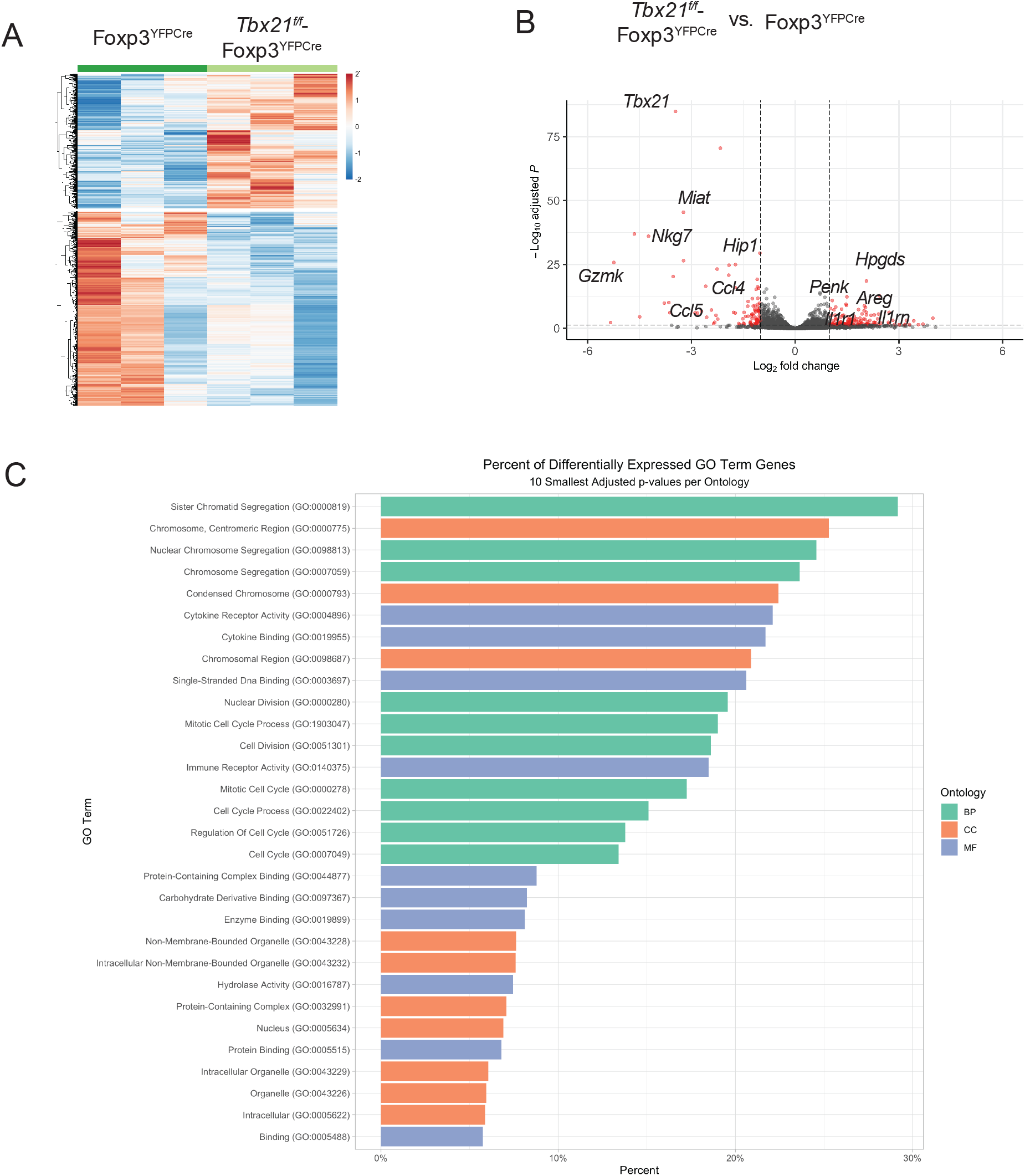
RNA-seq analysis shows that Tbet in Tregs governs a host of genes associated with fitness for Tregs during Th1 inflammation. Tregs were sorted from 9 d post-infected Foxp3^YFPCre^ and *Tbx21*^f/f^-Foxp3^YFPCre^ mice. RNA was isolated from the purified Tregs and RNA-Sequencing was performed. 3 mice of each genotype were combined per experiment and a total of three independent experiments were analyzed. (**A**) Heatmap of differentially expressed genes. (**B**) Volcano Plot highlighting differential genes in *Tbx21*^f/f^-Foxp3^YFPCre^ using Foxp3^YFPCre^ as reference. The most significantly regulated genes are in red and calculated using DESeq2. (**C**) GoSeq analysis of the top pathways in biological processes, cell cycle and metabolic function.

## Discussion

We show that the capacity to express Tbet in Tregs is necessary for survival of the host during acute *T. gondii* infection, as the majority of *Tbx21*^*f/f*^-Foxp3^YFPCre^, succumbed to the acute infection. Importantly, this increased susceptibility to infection was not due to uncontrolled parasite replication. These findings suggest that these Tbet-expressing Tregs are not required for regulating the protective immune response directed at the parasite. While there were minimal differences between the T helper cells found in the spleen, mesenteric lymph node and small intestinal lamina propria, mice harboring Tbet-deficient Tregs showed significantly reduced frequency and numbers of Tregs in the mesenteric lymph node and small intestinal lamina propria, respectively, in agreement with previous findings (5). We also observed that the frequency of Ki67 expression in Tregs was reduced in the small intestinal lamina propria of *Tbx21*^*f/f*^-Foxp3^YFPCre^ mice compared to Foxp3^YFPCre^ mice. Notably, serum IFN-γ and TNFα levels were significantly elevated in infected *Tbx21*^*f/f*^-Foxp3^YFPCre^ compared to Foxp3^YFPCre^ mice. These data present distinct differences when compared to global Tbet knockout mice, namely the loss of control of *T. gondii* replication and dissemination at peripheral sites during acute infection, and similar IFN-γ production compared to WT mice (39).

Moreover, it has been shown previously that Tbet-deficient Tregs have similar *in vitro* suppressive capacities WT Th1-Tregs (12, 40). Interestingly, in models of T cell mediated colitis, Tbet is not required for their suppressive capacity (21, 22, 40). This is in line with studies demonstrating that RoRγt-expressing Tregs are commensal specific and control intestinal inflammation (41, 42). Adding another layer to the understanding of Tbet-expressing Tregs, a recent study demonstrated that IFNγ^+^Tbet^+^ Tregs promoted Th-1 intestinal inflammation in the DSS colitis model (13). The differences between our findings and others could be explained by potential differences in the commensal flora between animal facilities, instigating inflammatory stimuli or the additional complexity of an ongoing infection in the gastrointestinal tract. Furthermore, Tbet is highly expressed in Tregs during *T. gondii* compared to other infections, which may alter its function in Tregs compared to Tregs expressing lower levels of Tbet (5, 6, 12, 14, 16, 19, 20). This is supported by data showing Tregs in a *T. gondii* infected environment are sensitive to IL-12 stimulation, resulting in Stat4 phosphorylation, while Tregs in a naive setting, or *L. monocytogenes*, are not IL-12 sensitive (6, 12). This sensitivity is Stat1-dependent and due to the sustained production of IL-12 and IL-27 in *T. gondii* infection (19). Prolonged Tbet expression in Tregs, however, appears to result in pathogenic Treg function and impair tissue remodeling programs during chronic infection of *T. gondii* of muscle (16–18). Together, our data shows distinct roles for Tregs during the acute response compared to the chronic response to *T. gondii* infection. It still remains to be fully explored how this prolonged Tbet expression may reprogram Tregs during later stages of infection.

The driving force of immune-mediated pathology in acute *T. gondii* infection is dysbiosis of the microbiota in the gastrointestinal tract (24, 37, 32, 35, 43–45). *Toxoplasma* infection can perturb the integrity of the epithelial barrier and cause a leaky gut (35, 37, 46, 47). This over-exuberant immune response is thought to be largely driven by CD4^+^ T cells (35, 37). In agreement with this, both depletion of CD4^+^ T cells and treatment with broad-spectrum antibiotics improved the disease course in *Tbx21*^*f/f*^-Foxp3^YFPCre^ compared to Foxp3^YFPCre^ mice. It is now well recognized that Tregs must adapt within the inflammatory environment in order to function and prevent this collateral damage. Their role in suppressing this damage from the parasite specific Th1-mediated immune response during *T. gondii* infection has been documented (48–50). Our study highlights a role for Tbet driving several aspects of fitness that promotes the ability of Tregs to dampen the immunopathology caused by dysbiosis during acute *T. gondii* infection. It remains to be determined the role of Tbet expression in Tregs during the chronic phase of infection in skeletal muscle and other tissues.

## Acknowledgements

Research reported in this publication was supported by the National Institute of Allergy and Infectious Diseases of the NIH under award number R21AI128284 (to EAW) and The Crohn’s and Colitis Foundation of America (ref #278937 to EAW). We thank the Confocal Imaging and Flow Cytometry Core, and the Genomics and Bioinformatics Core at the University at Buffalo, the National Institutes of Health Tetramer Core Facility for the *T. gondii* ME49 hypothetical protein tetramers, and Dr. Michael Grigg for generously providing the RFP-expressing ME49 parasite. We thank Dr. Joseph Barbi for his helpful discussion and critical reading of this article.

## References

1. Josefowicz, S. Z., L.-F. Lu, and A. Y. Rudensky. 2012. Regulatory T Cells: Mechanisms of Differentiation and Function. Annu. Rev. Immunol. 30: 531–564.

2. Kim, J. M., J. P. Rasmussen, and A. Y. Rudensky. 2007. Regulatory T cells prevent catastrophic autoimmunity throughout the lifespan of mice. Nat. Immunol. 8: 191–197.

3. Okeke, E. B., and J. E. Uzonna. 2019. The Pivotal Role of Regulatory T Cells in the Regulation of Innate Immune Cells. Front. Immunol. 10: 680.

4. Zheng, Y., A. Chaudhry, A. Kas, P. deRoos, J. M. Kim, T.-T. Chu, L. Corcoran, P. Treuting, U. Klein, and A. Y. Rudensky. 2009. Regulatory T-cell suppressor program co-opts transcription factor IRF4 to control T(H)2 responses. Nature 458: 351–356.

5. Koch, M. A., G. Tucker-Heard, N. R. Perdue, J. R. Killebrew, K. B. Urdahl, and D. J. Campbell. 2009. The transcription factor T-bet controls regulatory T cell homeostasis and function during type 1 inflammation. Nat. Immunol. 10: 595–602.

6. Koch, M. A., K. R. Thomas, N. R. Perdue, K. S. Smigiel, S. Srivastava, and D. J. Campbell. 2012. T-bet+ Treg Cells Undergo Abortive Th1 Cell Differentiation due to Impaired Expression of IL-12 Receptor β2. Immunity 37: 501–510.

7. Chaudhry, A., D. Rudra, P. Treuting, R. M. Samstein, Y. Liang, A. Kas, and A. Y. Rudensky. 2009. CD4+ regulatory T cells control TH17 responses in a Stat3-dependent manner. Science 326: 986–991.

8. Chung, Y., S. Tanaka, F. Chu, R. I. Nurieva, G. J. Martinez, S. Rawal, Y.-H. Wang, H. Lim, J. M. Reynolds, X. Zhou, H. Fan, Z. Liu, S. S. Neelapu, and C. Dong. 2011. Follicular regulatory T cells expressing Foxp3 and Bcl-6 suppress germinal center reactions. Nat. Med. 17: 983–988.

9. Wang, Y., M. A. Su, and Y. Y. Wan. 2011. An essential role of the transcription factor GATA-3 for the function of regulatory T cells. Immunity 35: 337–348.

10. Linterman, M. A., W. Pierson, S. K. Lee, A. Kallies, S. Kawamoto, T. F. Rayner, M. Srivastava, D. P. Divekar, L. Beaton, J. J. Hogan, S. Fagarasan, A. Liston, K. G. C. Smith, and C. G. Vinuesa. 2011. Foxp3+ follicular regulatory T cells control the germinal center response. Nat. Med. 17: 975–982.

11. Wohlfert, E. A., J. R. Grainger, N. Bouladoux, J. E. Konkel, G. Oldenhove, C. H. Ribeiro, J. A. Hall, R. Yagi, S. Naik, R. Bhairavabhotla, W. E. Paul, R. Bosselut, G. Wei, K. Zhao, M. Oukka, J. Zhu, and Y. Belkaid. 2011. GATA3 controls Foxp3^+^ regulatory T cell fate during inflammation in mice. J. Clin. Invest. 121: 4503–4515.

12. Oldenhove, G., N. Bouladoux, E. A. Wohlfert, J. A. Hall, D. Chou, L. Dos santos, S. O’Brien, R. Blank, E. Lamb, S. Natarajan, R. Kastenmayer, C. Hunter, M. E. Grigg, and Y. Belkaid. 2009. Decrease of Foxp3+ Treg Cell Number and Acquisition of Effector Cell Phenotype during Lethal Infection. Immunity 31: 772–786.

13. Di Giovangiulio, M., A. Rizzo, E. Franzè, F. Caprioli, F. Facciotti, S. Onali, A. Favale, C. Stolfi, H.-J. Fehling, G. Monteleone, and M. C. Fantini. 2019. Tbet Expression in Regulatory T Cells Is Required to Initiate Th1-Mediated Colitis. Front. Immunol. 10.

14. Levine, A. G., A. Mendoza, S. Hemmers, B. Moltedo, R. E. Niec, M. Schizas, B. E. Hoyos, E. V. Putintseva, A. Chaudhry, S. Dikiy, S. Fujisawa, D. M. Chudakov, P. M. Treuting, and A. Y. Rudensky. 2017. Stability and function of regulatory T cells expressing the transcription factor T-bet. Nature 546: 421.

15. Wohlfert, E., and Y. Belkaid. 2010. Plasticity of Treg at infected sites. Mucosal Immunol. 3: 213–215.

16. Jin, R. M., S. J. Blair, J. Warunek, R. R. Heffner, I. J. Blader, and E. A. Wohlfert. 2017. Regulatory T Cells Promote Myositis and Muscle Damage in Toxoplasma gondii Infection. J. Immunol. 198: 352–362.

17. Jin, R. M., J. Warunek, and E. A. Wohlfert. 2018. Therapeutic Administration of IL-10 and Amphiregulin Alleviates Chronic Skeletal Muscle Inflammation and Damage Induced by Infection. ImmunoHorizons 2: 142–154.

18. Jin, R. M., J. Warunek, and E. A. Wohlfert. 2018. Chronic infection stunts macrophage heterogeneity and disrupts immune-mediated myogenesis. JCI Insight 3.

19. Hall, A. O., D. P. Beiting, C. Tato, B. John, G. Oldenhove, C. G. Lombana, G. H. Pritchard, J. S. Silver, N. Bouladoux, J. S. Stumhofer, T. H. Harris, J. Grainger, E. D. T. Wojno, S. Wagage, D. S. Roos, P. Scott, L. A. Turka, S. Cherry, S. L. Reiner, D. Cua, Y. Belkaid, M. M. Elloso, and C. A. Hunter. 2012. The Cytokines Interleukin 27 and Interferon-γ Promote Distinct Treg Cell Populations Required to Limit Infection-Induced Pathology. Immunity 37: 511–523.

20. O’Brien, C. A., C. Overall, C. Konradt, A. C. O. Hall, N. W. Hayes, S. Wagage, B. John, D. A. Christian, C. A. Hunter, and T. H. Harris. 2017. CD11c-Expressing Cells Affect Regulatory T Cell Behavior in the Meninges during Central Nervous System Infection. J. Immunol. 198: 4054–4061.

21. McPherson, R. C., D. G. Turner, I. Mair, R. A. O’Connor, and S. M. Anderton. 2015. T-bet Expression by Foxp3+ T Regulatory Cells is Not Essential for Their Suppressive Function in CNS Autoimmune Disease or Colitis. Front. Immunol. 6.

22. Yu, F., S. Sharma, J. Edwards, L. Feigenbaum, and J. Zhu. 2015. Dynamic expression of transcription factors T-bet and GATA-3 by regulatory T cells maintains immunotolerance. Nat. Immunol. 16: 197–206.

23. Rakoff-Nahoum, S., J. Paglino, F. Eslami-Varzaneh, S. Edberg, and R. Medzhitov. 2004. Recognition of Commensal Microflora by Toll-Like Receptors Is Required for Intestinal Homeostasis. Cell 118: 229–241.

24. Molloy, M. J., J. R. Grainger, N. Bouladoux, T. W. Hand, L. Y. Koo, S. Naik, M. Quinones, A. K. Dzutsev, J.-L. Gao, G. Trinchieri, P. M. Murphy, and Y. Belkaid. 2013. Intraluminal containment of commensal outgrowth in the gut during infection-induced dysbiosis. Cell Host Microbe 14: 318–328.

25. Suzuki, Y., M. A. Orellana, R. D. Schreiber, and J. S. Remington. 1988. Interferon-gamma: the major mediator of resistance against Toxoplasma gondii. Science 240: 516–518.

26. Hunter, C. A., C. S. Subauste, V. H. Van Cleave, and J. S. Remington. 1994. Production of gamma interferon by natural killer cells from Toxoplasma gondii-infected SCID mice: regulation by interleukin-10, interleukin-12, and tumor necrosis factor alpha. Infect. Immun. 62: 2818–2824.

27. Deckert-Schlüter, M., H. Bluethmann, A. Rang, H. Hof, and D. Schlüter. 1998. Crucial Role of TNF Receptor Type 1 (p55), But Not of TNF Receptor Type 2 (p75), in Murine Toxoplasmosis. J. Immunol. 160: 3427–3436.

28. Yap, G. S., T. Scharton-Kersten, H. Charest, and A. Sher. 1998. Decreased Resistance of TNF Receptor p55- and p75-Deficient Mice to Chronic Toxoplasmosis Despite Normal Activation of Inducible Nitric Oxide Synthase In Vivo. J. Immunol. 160: 1340–1345.

29. Schlüter, D., L.-Y. Kwok, S. Lütjen, S. Soltek, S. Hoffmann, H. Körner, and M. Deckert. 2003. Both Lymphotoxin-α and TNF Are Crucial for Control of Toxoplasma gondii in the Central Nervous System. J. Immunol. 170: 6172–6182.

30. Gazzinelli, R. T., M. Wysocka, S. Hieny, T. Scharton-Kersten, A. Cheever, R. Kühn, W. Müller, G. Trinchieri, and A. Sher. 1996. In the absence of endogenous IL-10, mice acutely infected with Toxoplasma gondii succumb to a lethal immune response dependent on CD4+ T cells and accompanied by overproduction of IL-12, IFN-gamma and TNF-alpha. J. Immunol. 157: 798–805.

31. Villarino, A., L. Hibbert, L. Lieberman, E. Wilson, T. Mak, H. Yoshida, R. A. Kastelein, C. Saris, and C. A. Hunter. 2003. The IL-27R (WSX-1) Is Required to Suppress T Cell Hyperactivity during Infection. Immunity 19: 645–655.

32. Liesenfeld, O., J. Kosek, J. S. Remington, and Y. Suzuki. 1996. Association of CD4+ T cell-dependent, interferon-gamma-mediated necrosis of the small intestine with genetic susceptibility of mice to peroral infection with Toxoplasma gondii. J. Exp. Med. 184: 597–607.

33. Liesenfeld, O. 2002. Oral infection of C57BL/6 mice with Toxoplasma gondii: a new model of inflammatory bowel disease? J. Infect. Dis. 185 Suppl 1: S96–101.

34. Egan, C. E., S. B. Cohen, and E. Y. Denkers. 2012. Insights into inflammatory bowel disease using Toxoplasma gondii as an infectious trigger. Immunol. Cell Biol. 90: 668–675.

35. Raetz, M., S. Hwang, C. Wilhelm, D. Kirkland, A. Benson, C. Sturge, J. Mirpuri, S. Vaishnava, B. Hou, A. L. DeFranco, C. J. Gilpin, L. V. Hooper, and F. Yarovinsky. 2013. Parasite-induced TH1 cells and intestinal dysbiosis cooperate in IFN-γ-dependent elimination of Paneth cells. Nat. Immunol. 14: 136–142.

36. Burger, E., A. Araujo, A. López-Yglesias, M. W. Rajala, L. Geng, B. Levine, L. V. Hooper, E. Burstein, and F. Yarovinsky. 2018. Loss of Paneth Cell Autophagy Causes Acute Susceptibility to Toxoplasma gondii-Mediated Inflammation. Cell Host Microbe 23: 177-190.e4.

37. Heimesaat, M. M., S. Bereswill, A. Fischer, D. Fuchs, D. Struck, J. Niebergall, H.-K. Jahn, I. R. Dunay, A. Moter, D. M. Gescher, R. R. Schumann, U. B. Göbel, and O. Liesenfeld. 2006. Gram-Negative Bacteria Aggravate Murine Small Intestinal Th1-Type Immunopathology following Oral Infection with Toxoplasma gondii. J. Immunol. 177: 8785–8795.

38. Craven, M., C. E. Egan, S. E. Dowd, S. P. McDonough, B. Dogan, E. Y. Denkers, D. Bowman, E. J. Scherl, and K. W. Simpson. 2012. Inflammation Drives Dysbiosis and Bacterial Invasion in Murine Models of Ileal Crohn’s Disease. PLoS ONE 7.

39. Pritchard, G. H., A. O. Hall, D. A. Christian, S. Wagage, Q. Fang, G. Muallem, B. John, A. G. Zaretsky, W. G. Dunn, J. Perrigoue, S. L. Reiner, and C. A. Hunter. 2015. Diverse Roles for T-bet in the Effector Responses Required for Resistance to Infection. J. Immunol. 194: 1131–1140.

40. Feng, T., A. T. Cao, C. T. Weaver, C. O. Elson, and Y. Cong. 2011. Interleukin-12 Converts Foxp3+ Regulatory T Cells to Interferon–γ-Producing Foxp3+ T Cells That Inhibit Colitis. Gastroenterology 140: 2031–2043.

41. Ohnmacht, C., J.-H. Park, S. Cording, J. B. Wing, K. Atarashi, Y. Obata, V. Gaboriau-Routhiau, R. Marques, S. Dulauroy, M. Fedoseeva, M. Busslinger, N. Cerf-Bensussan, I. G. Boneca, D. Voehringer, K. Hase, K. Honda, S. Sakaguchi, and G. Eberl. 2015. MUCOSAL IMMUNOLOGY. The microbiota regulates type 2 immunity through RORγt^+^ T cells. Science 349: 989–993.

42. Sefik, E., N. Geva-Zatorsky, S. Oh, L. Konnikova, D. Zemmour, A. M. McGuire, D. Burzyn, A. Ortiz-Lopez, M. Lobera, J. Yang, S. Ghosh, A. Earl, S. B. Snapper, R. Jupp, D. Kasper, D. Mathis, and C. Benoist. 2015. Individual intestinal symbionts induce a distinct population of RORγ+ regulatory T cells. Science 349: 993–997.

43. Bereswill, S., M. Muñoz, A. Fischer, R. Plickert, L.-M. Haag, B. Otto, A. A. Kühl, C. Loddenkemper, U. B. Göbel, and M. M. Heimesaat. 2010. Anti-Inflammatory Effects of Resveratrol, Curcumin and Simvastatin in Acute Small Intestinal Inflammation. PLoS ONE 5.

44. Bereswill, S., A. A. Kühl, M. Alutis, A. Fischer, L. Möhle, D. Struck, O. Liesenfeld, U. B. Göbel, I. R. Dunay, and M. M. Heimesaat. 2014. The impact of Toll-like-receptor-9 on intestinal microbiota composition and extra-intestinal sequelae in experimental Toxoplasma gondii induced ileitis. Gut Pathog. 6: 19.

45. Heimesaat, M. M., I. R. Dunay, M. Alutis, A. Fischer, L. Möhle, U. B. Göbel, A. A. Kühl, and S. Bereswill. 2014. Nucleotide-Oligomerization-Domain-2 Affects Commensal Gut Microbiota Composition and Intracerebral Immunopathology in Acute Toxoplasma gondii Induced Murine Ileitis. PLOS ONE 9: e105120.

46. McLeod, R., P. Eisenhauer, D. Mack, C. Brown, G. Filice, and G. Spitalny. 1989. Immune responses associated with early survival after peroral infection with Toxoplasma gondii. J. Immunol. 142: 3247–3255.

47. Schreiner, M., and O. Liesenfeld. 2009. Small intestinal inflammation following oral infection with Toxoplasma gondii does not occur exclusively in C57BL/6 mice: review of 70 reports from the literature. Mem. Inst. Oswaldo Cruz 104: 221–233.

48. Morampudi, V., S. De Craeye, A. Le Moine, S. Detienne, M. Y. Braun, and S. D’Souza. 2011. Partial depletion of CD4+CD25+Foxp3+ T regulatory cells significantly increases morbidity during acute phase Toxoplasma gondii infection in resistant BALB/c mice. Microbes Infect. 13: 394–404.

49. Tenorio, E. P., J. E. Olguín, J. Fernández, P. Vieyra, and R. Saavedra. 2010. Reduction of Foxp3+ Cells by Depletion with the PC61 mAb Induces Mortality in Resistant BALB/c Mice Infected with Toxoplasma gondii. J. Biomed. Biotechnol. 2010.

50. Couper, K. N., P. A. Lanthier, G. Perona-Wright, L. W. Kummer, W. Chen, S. T. Smiley, M. Mohrs, and L. L. Johnson. 2009. Anti-CD25 Antibody-Mediated Depletion of Effector T Cell Populations Enhances Susceptibility of Mice to Acute but Not Chronic Toxoplasma gondii Infection. J. Immunol. 182: 3985–3994.

